# PPP1R3B is a metabolic switch that shifts hepatic energy storage from lipid to glycogen

**DOI:** 10.1101/2023.03.04.529958

**Authors:** Kate Townsend Creasy, Minal B. Mehta, Joseph Park, Carolin V. Schneider, Swapnil V. Shewale, John S. Millar, Nicholas J. Hand, Joseph A. Baur, Daniel J. Rader

## Abstract

Obesity is a growing worldwide epidemic that carries numerous metabolic complications including increased risk of type 2 diabetes (T2D), cardiovascular disease (CVD), and non-alcoholic fatty liver disease (NAFLD). Multiple genome-wide association studies (GWAS) have associated the *PPP1R3B* locus with cardiometabolic traits including fasting glucose and insulin levels (T2D traits), plasma lipids (CVD traits), and indications of hepatic steatosis and liver damage (NAFLD traits)^1–5^. The *PPP1R3B* gene encodes the glycogen regulatory protein PPP1R3B (also known as G_L_) which has an established role in liver glycogen metabolism and plasma glucose homeostasis^6,7^. The metabolic and NAFLD GWAS single nucleotide polymorphisms (SNPs) in this region, which are all in high linkage disequilibrium, result in increased liver *PPP1R3B* expression and hepatic glycogen accumulation, but have provided conflicting results on the impacts on hepatic steatosis and liver damage. Here we investigate the consequences of both *Ppp1r3b* overexpression and deletion in mouse and cell models and find that dysregulated *Ppp1r3b* expression in either direction promotes metabolic dysfunction and liver injury. Hepatocyte overexpression of *Ppp1r3b* increases hepatic glycogen storage, prolongs fasting blood glucose levels, and confers protection from hepatic steatosis, but increases plasma ALT in aged animals. Conversely, deletion of hepatocyte *Ppp1r3b* eliminates hepatic glycogen, causes impaired glucose disposal, and results in hepatic steatosis with age or high sucrose diet. We investigated the metabolic pathways contributing to steatosis and found that *Ppp1r3b* deletion and diminished glycogenesis diverts the storage of exogenous glucose to hepatic triglycerides (TG), and stored liver lipids are preferentially used for energy during fasting through lipid oxidation and ketogenesis. Further, we interrogated two large human biobank cohorts and found carriers of SNPs associated with increased *PPP1R3B* expression have increased plasma glucose, decreased hepatic fat, and lower plasma lipids, while putative loss-of-function (pLoF) variant carriers have increased hepatic fat and elevated plasma ketones and lipids, consistent with the results seen in our mouse models. These findings suggest hepatic PPP1R3B serves as a metabolic switch favoring hepatic energy storage as glycogen instead of TG.

## Main

NAFLD is is one of the fastest growing metabolic diseases in the world, exacerbated by the obesity epidemic and increasing prevalence of T2D and other metabolic disorders^8^. NALFD is a spectrum of liver conditions that ranges from the initial accumulation of excess TG in lipid droplets in the liver (steatosis), advancing to inflammatory steatohepatitis (NASH), and further to fibrosis, cirrhosis, and hepatocellular carcinoma (HCC), with diminishing liver function and increasing risk of comorbidities with disease progression. Advanced stages of NAFLD are a growing public health concern and are currently among the leading indications for liver transplant due to failure^9^. NAFLD is highly prevalent among patients with obesity (60-95%), T2D (55-70%), and Metabolic Syndrome (43-67%), and is also highly heritable^10,11^. Therefore, investigating the mechanisms and metabolic consequences of genes associated with NAFLD and related cardiometabolic disorders may help elucidate the pathophysiology of these disorders to develop better screening and diagnostic criteria as well as therapeutic strategies.

Several GWAS have associated a chromosomal region at 8p23.1, named *PPP1R3B* for the nearest coding gene, with a number of metabolic traits including elevated fasting glucose, insulin, and lactate levels^1,12^, lower plasma lipids^13^, as well as increased liver enzymes ALT and ALP^2,5,14^. The variants at these GWAS signals comprise a tight linkage block with the sentinel SNPs from all relevant GWAS falling within 873bp of one another and overlapping the proximal promoter and 5’ end of a long non-coding RNA (lncRNA) *LOC157273*. The lead SNP associated with lower plasma lipids, rs9987289, was also reported to be a liver expression quantitative trait locus (eQTL) for *PPP1R3B*, with the minor allele associated with higher liver *PPP1R3B* transcript levels^13^. Because the minor allele is associated with higher fasting glucose, lower plasma lipids, and higher ALT/ALP, this suggested that increased hepatic *PPP1R3B* expression confers these phenotypes. Recent studies support that the *LOC157273* lncRNA is a transcriptional repressor of *PPP1R3B* expression, as carriers of the minor allele have lower abundance of this lncRNA and increased *PPP1R3B* expression, while CRISPR/Cas9 deletion in HuH7 cells leads to decreased *LOC157273* mRNA and increased *PPP1R3B* expression^15,16^.

The *PPP1R3B* gene encodes the glycogen regulatory protein <u>p</u>rotein <u>p</u>hos<u>p</u>hatase <u>1</u> (PP1) regulatory subunit <u>3B</u> (historically known as G_L_). PPP1R3B recruits PP1 to post-translationally regulate, through dephosphorylation, the increased activity of Glycogen Synthase (GS) and decreased activity of Glycogen Phosphorylase (GP), the rate-limiting enzymes in glycogen synthesis and hydrolysis, respectively, which effectively increases liver glycogen stores^6,17^ (**Fig. 1a**). The glycemic and T2D GWAS associations are concordant with the functional roles of PPP1R3B in glycogen metabolism and glucose homeostasis that have been demonstrated in mouse and cell models^7,18^. However, it remains unclear how *PPP1R3B* expression is associated with hepatic and plasma lipid metabolism as suggested by GWAS. The *PPP1R3B/LOC157273* locus was identified as a signal for hepatic steatosis by computed tomography (CT), although there was no histological evidence of steatosis in matched liver biopsies^2,19^. Subsequent GWAS and mechanistic studies have since provided strong evidence that the *LOC157273* SNPs (with increased *PPP1R3B* expression) are actually a signal for increased hepatic glycogen, which alters liver attenuation and can be ambiguous on liver imaging, and that increased glycogen storage contributes to mild liver damage that is reflected in elevated plasma ALT^15,18,20,21^. In fact, studies in several European population cohorts support that the reported *LOC157273* SNPs have less hepatic lipid accumulation assessed by imaging and that *LOC157273* variant carriers with traditional NAFLD risk factors were protected from steatosis^18,20,22^. We recently reported concurring findings that the *LOC157273* locus is associated with chronically elevated ALT, but less hepatic fat estimated by radiological imaging^5^. *PPP1R3B* has been suggested as a T2D therapeutic target, as *PPP1R3B* overexpression and increased hepatic glycogen synthesis improves postprandial glucose clearance from the blood^23,24^, and thus the consequences of altered *PPP1R3B* expression on liver damage and other metabolic phenotypes should be thoroughly examined.

**Figure 1.**
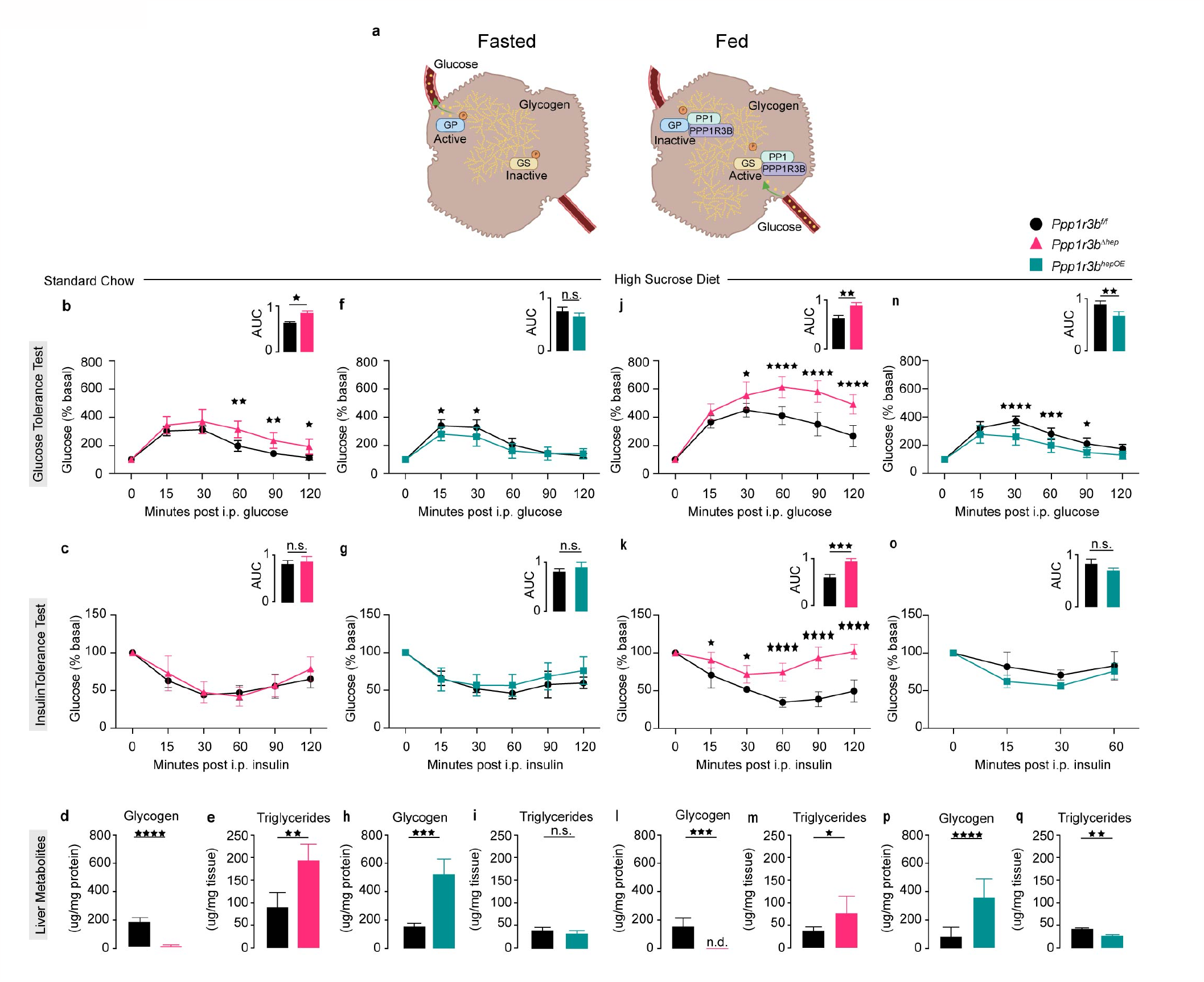
*Ppp1r3b*^*∆hep*^ mice develop glucose intolerance and steatosis with aging or sucrose diet challenge, while *Ppp1r3b*^*OE*^ mice have increased liver glycogen and reduced triglyceride content. **a**. Illustration of the function of PPP1R3B in hepatocytes: PPP1R3B is a scaffolding protein that facilitates the physical interaction of Protein Phosphatase 1 (PP1) to targets Glycogen Synthase (GS) and Glycogen Phosphorylase (GP), thereby regulating the activity status of glycogen metabolizing enzymes and postprandial glucose metabolism. **b-e:** Metabolic tests performed on fasted 6-9 months aged *Ppp1r3b*^*f/f*^ (black, n=6) and *Ppp1r3b*^*∆hep*^ (pink, n=8) mice maintained on chow diet. **b**. Intraperitoneal glucose tolerance test (area under the curve (AUC) inset). **c**. Insulin tolerance test, AUC inset. **d**. Hepatic glycogen content. **e**. hepatic TG content. **f-i:** Metabolic tests performed on fasted 6-9 months aged *Ppp1r3b*^*f/f*^ (black, n=8) and *Ppp1r3b*^*hepOE*^ (green, n=6) mice maintained on chow diet. **f**. Intraperitoneal glucose tolerance test, AUC inset. **g**. Insulin tolerance test, AUC inset. **h**. Hepatic glycogen content. **i**. hepatic TG content. **j-m:** Metabolic tests performed on fasted *Ppp1r3b*^*f/f*^ (black, n=5) and *Ppp1r3b*^*∆hep*^ (pink, n=7) mice on High Sucrose Diet (65%) for 12 weeks. **j**. Intraperitoneal glucose tolerance test, AUC inset. **k**. Insulin tolerance test, AUC inset. **l**. Hepatic glycogen content. **m**. hepatic TG content. **n-q:** Metabolic tests performed on fasted *Ppp1r3b*^*f/f*^ (black, n=8) and *Ppp1r3b*^*hepOE*^ (green, n=8) on High Sucrose Diet for 12 weeks. **n**. Intraperitoneal glucose tolerance test, AUC inset. **o**. Insulin tolerance test, AUC inset. **p**. Hepatic glycogen content. **q**. Hepatic TG content.

As *LOC157273* is expressed in humans and non-human primates but is not conserved in rodents^15^, we investigated the metabolic consequences of altered *Ppp1r3b* expression by directly targeting *Ppp1r3b* expression in mice. We previously showed that perturbing hepatocyte *Ppp1r3b* expression in genetically modified mice impacts glycogen metabolism and glycemic traits: hepatocyte-specific deletion of *Ppp1r3b* (*Ppp1r3b*^*∆hep*^) causes dramatic reduction in glycogen synthase activity, depletion of liver glycogen stores, and rapid fasting hypoglycemia, whereas hepatocyte overexpression of murine *Ppp1r3b* (*Ppp1r3b*^*hepOE*^) increases liver glycogen content and preserves blood glucose levels even after prolonged fasting compared to wildtype mice *(Ppp1r3b*^*f/f*^)^7^. We examined the metabolic consequences of prolonged dysglycemia in aged (6-9 months) mice maintained on chow and found that aged *Ppp1r3b*^*∆hep*^ mice have impaired glucose disposal assessed by glucose tolerance test (GTT), even while maintaining normal insulin response during insulin tolerance test (ITT) (**Fig. 1b-c**). *Ppp1r3b*^*∆hep*^ mice had severely diminished hepatic glycogen levels as expected (*Ppp1r3b*^*f/f*^: 178.2 µg/mg protein; *Ppp1r3b*^*∆hep*^: 2.4µg/mg; **Fig. 1d**), and we observed increased liver TG in the aged *Ppp1r3b*^*∆hep*^ mice compared to *Ppp1r3b*^*f/f*^ controls (*Ppp1r3b*^*f/f*^: 90.0 µg/mg tissue mass; *Ppp1r3b*^*∆hep*^: 193.7 µg/mg; **Fig. 1e**). The reciprocal phenotype appears in aged *Ppp1r3b*^*hepOE*^ mice, as they have improved glucose disposal during GTT (**Fig. 1f**) and normal insulin response (**Fig. 1g**). *Ppp1r3b*^*hepOE*^ mice maintained increased hepatic glycogen stores (*Ppp1r3b*^*f/f*^: 154.2 µg/mg; *Ppp1r3b*^*hepOE*^: 523.5 µg/mg; **Fig.1h**) without differences in liver TG compared to *Ppp1r3b*^*f/f*^ (*Ppp1r3b*^*f/f*^: 30.1 µg/mg; *Ppp1r3b*^*hepOE*^: 21.6 µg/mg; **Fig. 1i**). As PPP1R3B has a critical role in glucose metabolism and glycemic traits, we next evaluated the metabolic responses to changes in hepatic *Ppp1r3b* expression in mice challenged with a high sucrose diet (HSD, 66% sucrose) for 12 weeks. The *Ppp1r3b*^*∆hep*^ mice developed dramatically impaired glucose tolerance and insulin insensitivity compared to *Ppp1r3b*^*f/f*^ mice, assessed by GTT and ITT (**Fig. 1j-k**). Conversely, the *Ppp1r3b*^*hepOE*^ mice continued to display improved glucose disposal and normal insulin response (**Fig. 1n-o**). Again, the *Ppp1r3b*^*∆hep*^ mice had negligible liver glycogen (**Fig. 1l**) but increased hepatic TG compared to *Ppp1r3b*^*f/f*^ mice (*Ppp1r3b*^*f/f*^: 34.7 µg/mg protein; *Ppp1r3b*^*∆hep*^: 80.2 µg/mg, **Fig. 1m**), while *Ppp1r3b*^*hepOE*^ mice had increased hepatic glycogen (*Ppp1r3b*^*f/f*^: 84.1 µg/mg; *Ppp1r3b*^*hepOE*^: 363.9 µg/mg) and reduced hepatic TG compared to *Ppp1r3b*^*f/f*^ (*Ppp1r3b*^*f/f*^: 31.4 µg/mg; *Ppp1r3b*^*hepOE*^: 19.9 µg/mg; **Fig. 1p-q**). Overexpression of hepatic *Ppp1r3b* in mice was protective from dysglycemia and hepatic lipid accumulation with age and carbohydrate dietary challenge, which is consistent with the human GWAS findings and studies in NAFLD-risk populations. Interestingly, *Ppp1r3b*^*∆hep*^ mice had elevated hepatic TG content both in aged mice on chow and with HSD, which has not been previously reported in either human studies or other mouse models of altered *Ppp1r3b* expression.

In the postprandial state, hepatic glucose is rapidly converted to glucose-6-phosphate which then acts as an allosteric activator of GS to increase glycogen stores after fasting. After prolonged fasting when glycogen stores are depleted, hepatocytes utilize TG as the primary source of ATP^25^. We previously demonstrated that ablation of *Ppp1r3b* in mouse hepatocytes diminishes the conversion of postprandial glucose into hepatic glycogen, causing lower fasting plasma glucose and accelerated activation of gluconeogenic genes^7^. Since defective glucose-to-glycogen synthesis and increased gluconeogenesis are associated with hepatic steatosis, we investigated the impact of *Ppp1r3b* expression on hepatic TG accumulation. Littermate 8-weeks-old *Ppp1r3b*^*f/f*^ mice were administered hepatocyte-specific (TBG promoter) adeno-associated virus (AAV) with either *Cre recombinase* (*Ppp1r3b*^*∆hep*^), mouse *Ppp1r3b* (*Ppp1r3b*^*hepOE*^), or empty expression cassette as control (*Ppp1r3b*^*f/f*^) and were maintained on chow. At 4-months of age, *Ppp1r3b*^*∆hep*^ mice had significantly reduced 4-hours fasting blood glucose compared to both *Ppp1r3b*^*f/f*^ and *Ppp1r3b*^*hepOE*^ mice (*Ppp1r3b*^*f/f*^: 145.2 mg/dL; *Ppp1r3b*^*∆hep*^: 77.9 mg/dL; *Ppp1r3b*^*hepOE*^: 168 mg/dL; **Fig. 2a**), while there is no difference in fasting insulin or lactate levels (**Fig. 2b-c**). *Ppp1r3b*^*∆hep*^ mice had elevated fasting blood ketones (*Ppp1r3b*^*f/f*^: 0.46 mmol/L; *Ppp1r3b*^*∆hep*^: 1.05 mmol/L; *Ppp1r3b*^*hepOE*^: 0.6 mmol/L; **Fig. 2d**), indicative of lipid oxidation as an alternate fasting energy source. Upon sacrifice, *Ppp1r3b*^*hepOE*^ mice had an increased liver/body weight ratio (**Fig. 2e**), reflecting increased liver glycogen stores (**Fig. 2f**) and reduced liver TG (**Fig. 2g**). *Ppp1r3b*^*∆hep*^ mice had increased liver TG compared to both *Ppp1r3b*^*f/f*^ and *Ppp1r3b*^*hepOE*^ mice (**Fig. 2g**). There was also histological evidence of steatosis in livers from *Ppp1r3b*^*∆hep*^ mice marked by positive immunohistochemical staining for the lipid droplet-associated protein PLIN2 (**Fig. 2k**), while *Ppp1r3b*^*hepOE*^ livers appeared free of lipid droplets and had increased glycogen deposition visible with Periodic acid-Schiff’s (PAS) staining (**Fig. 2j-k**). Interestingly, both the *Ppp1r3b*^*∆hep*^ and *Ppp1r3b*^*hepOE*^ mice had elevated plasma ALT compared to *Ppp1r3b*^*f/f*^ mice (*Ppp1r3b*^*f/f*^: 44.2 U/L; *Ppp1r3b*^*∆hep*^: 71.2 U/L; *Ppp1r3b*^*hepOE*^: 79.2 U/L; **Fig. 2h**), suggesting that both deletion and overexpression of hepatocyte *Ppp1r3b* contribute to liver damage, with *Ppp1r3b*^*∆hep*^ mice accumulating more hepatic TG and *Ppp1r3b*^*hepOE*^ mice storing more glycogen.

**Figure 2.**
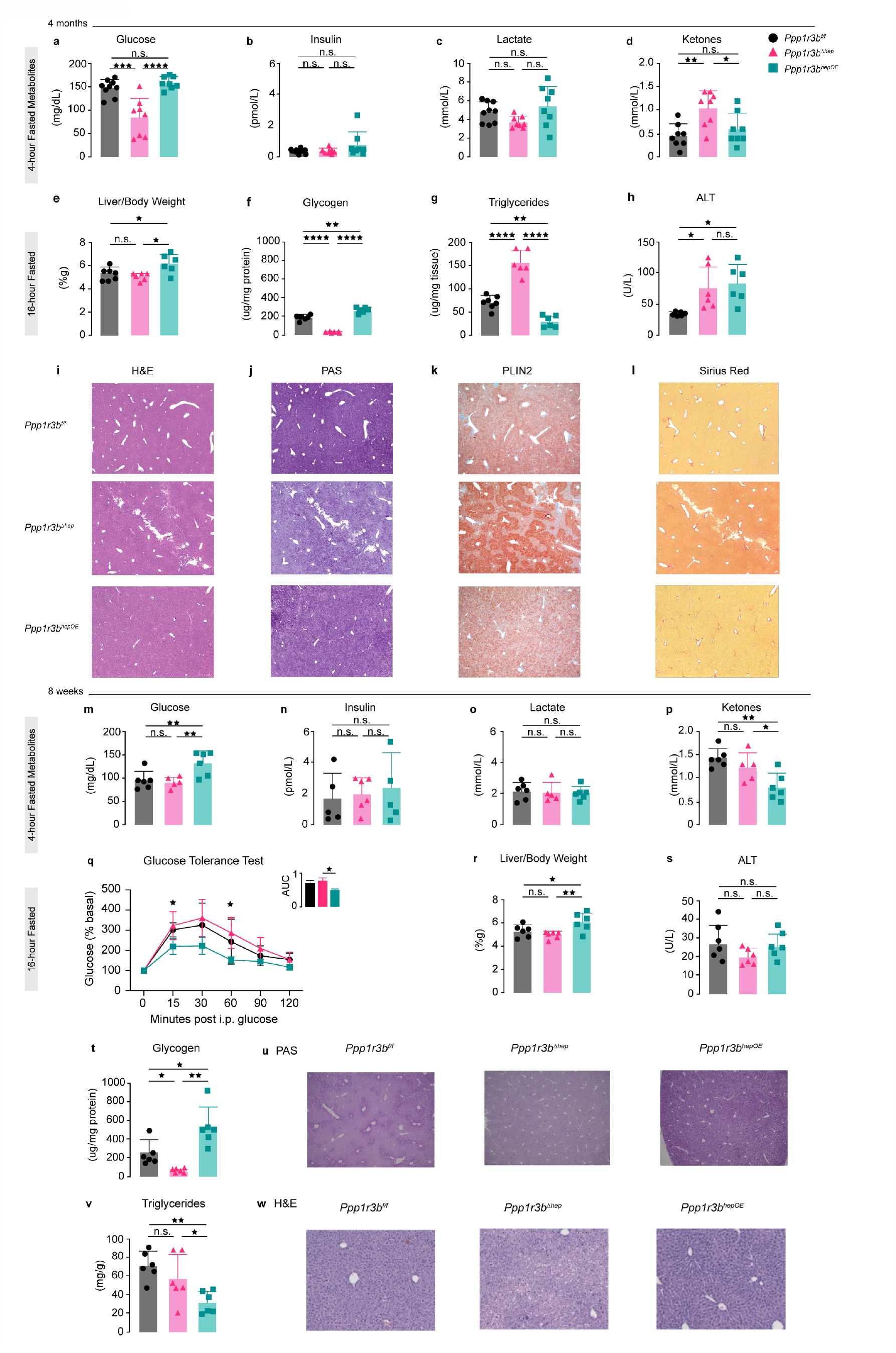
Mice with hepatocyte *Ppp1r3b* deletion develop metabolic dysfunction contributing to hepatic steatosis with age, while *Ppp1r3b* overexpressing mice rapidly increase glycogen synthesis and storage contributing to liver damage with aging. <u>Top</u>: Metabolic phenotypes of 4-months aged mice. **a-d**: Four-hour fasted metabolites in *Ppp1r3b*^*f/f*^ (black, n=9), *Ppp1r3b*^*∆hep*^ (pink, n=8), and *Ppp1r3b*^*hepOE*^ (green, n=8) mice. **a**. Blood glucose. **b**. Plasma insulin. **c**. Blood lactate. **d**. Blood ketones. **e-l:** Characteristics of overnight (16 hours) fasted *Ppp1r3b*^*f/f*^ (black, n=7), *Ppp1r3b*^*∆hep*^ (pink, n=6), and *Ppp1r3b*^*hepOE*^ (green, n=6) mice. **e**. Liver-to-body weight ratio. **f**. Hepatic glycogen content. **g**. Hepatic TG content. **h**. Fasted plasma ALT measurements. **i-l:** Paraffin embedded liver sections sectioned and stained with **i**. hematoxylin and eosin (H&E), **j**. Periodic acid-Schiff’s stain (PAS) to detect polysaccharide content, **k**. immunohistochemistry with PLIN2 antibodies to indicate lipid droplets, and **l**. Picro-Sirius Red to detect collagen deposition and liver lesions. <u>Bottom</u>: Metabolic phenotypes of 8-weeks old *Ppp1r3b*^*f/f*^ (black, n=6), *Ppp1r3b*^*∆hep*^ (pink, n=5), and *Ppp1r3b*^*hepOE*^ (green, n=6) mice. Four-hour fasted blood **m**. glucose, **n**. insulin, **o**. lactate, and **p**. ketones. **q-v:** Characteristics of overnight (16 hours) fasted *Ppp1r3b*^*f/f*^ (black, n=6), *Ppp1r3b*^*∆hep*^ (pink, n=6), and *Ppp1r3b*^*hepOE*^ (green, n=6) mice. **q**. Intraperitoneal glucose tolerance test, AUC inset. **r**. Liver-to-body weight ratio. **s**. Fasted plasma ALT measurements. **t**. Hepatic glycogen content. **u**. PAS staining of paraffin embedded liver sections. **v**. Hepatic TG content. **w**. PLIN2 immunohistochemistry. Group average presented as bar + s.d. with individual animals represented as shapes.

Stendler *et al*. reported that adenovirus mediated hepatic *Ppp1r3b* overexpression increased liver glycogen while constitutive *Ppp1r3b* deletion caused significant liver glycogen depletion, but there was no change in hepatic TG levels from either *Ppp1r3b* deletion or overexpression^18^. These studies were performed in young chow-fed mice, so we next characterized the metabolic phenotypes in young chow fed mice looking specifically at the impact of altered hepatocyte *Ppp1r3b* expression on molecular metabolism that could explain the observed steatosis phenotypes in older and high sucrose fed mice. Littermate *Ppp1r3b*^*f/f*^ mice were administered AAVs at five weeks of age and examined after three weeks (at 8-weeks old) and assessed for fasted circulating metabolites. After 4 hours fasting, *Ppp1r3b*^*hepOE*^ mice maintained higher blood glucose and both *Ppp1r3b*^*f/f*^ and *Ppp1r3b*^*∆hep*^ mice (Fig. 2m). There were no differences in fasting insulin or lactate (**Fig. 2n-o**), but *Ppp1r3b*^*hepOE*^ mice had significantly reduced fasting blood ketones than *Ppp1r3b*^*f/f*^ and *Ppp1r3b*^*∆hep*^ mice (*Ppp1r3b*^*f/f*^: 1.4 mmol/L; *Ppp1r3b*^*∆hep*^: 1.2 mmol/L; *Ppp1r3b*^*hepOE*^: 0.8 mmol/L; Fig. 2p). When examined by GTT after an overnight fast, the *Ppp1r3b*^*hepOE*^ mice had significantly improved glucose disposal, while there was no difference in *Ppp1r3b*^*∆hep*^ mice compared to *Ppp1r3b*^*f/f*^ (**Fig. 2q**). Mice were allowed to recover from the GTT and were sacrificed at 9-weeks old after an overnight fast. Examination of the livers revealed that *Ppp1r3b*^*hepOE*^ mice had increased liver-to-body weight ratio (**Fig. 2r**) and a corresponding increase in glycogen stores (**Fig. 2t-u**). Remarkably, livers from *Ppp1r3b*^*hepOE*^ mice had significantly less hepatic TG by both biochemical measurements (*Ppp1r3b*^*f/f*^: 71.8mg/g; *Ppp1r3b*^*∆hep*^: 58.1mg/g; *Ppp1r3b*^*hepOE*^: 32.3mg/g, p<0.01; **Fig. 2v**) and histologically (**Fig. 2w**) after only three weeks of induced *Ppp1r3b* overexpression. The 8-weeks old *Ppp1r3b*^*∆hep*^ mice did not display any differences in liver TG compared to *Ppp1r3b*^*f/f*^ mice, consistent with what has been previously reported^18^. A similar phenotype was reported in mice with liver-specific deletion of GS (gene *Gys2*) with greatly diminished liver glycogenesis in which knockout mice have increased liver TG compared to controls at 7-months and 15-months of age, but not earlier timepoints^26,27^. These data demonstrate that the metabolic changes and liver phenotypes associated with overexpression of *Ppp1r3b* occur very rapidly while hepatic lipid accumulation in *Ppp1r3b*^*∆hep*^ manifests over time was only observed in mice 4-months or older, or after HSD challenge.

To elucidate the mechanisms underlying the opposing effects on steatosis we observed in *Ppp1r3b*^*∆hep*^ and *Ppp1r3b*^*hepOE*^ mice, we investigated four common pathways that contribute to an imbalance in hepatic lipid accumulation and utilization that are often dysregulated in hepatic steatosis: 1) increased uptake and re-esterification of adipose-derived non-esterified fatty acids (NEFA); 2) increased *de novo* lipogenesis (DNL); 3) decreased TG secretion in VLDL particles; and 4) decreased β-oxidation of lipids. Low blood glucose levels and the corresponding decrease in circulating insulin in acute fasting elicits a cascade of signaling responses to the liver to produce ATP from available sources including glycogenolysis of hepatic glycogen stores, the use of lactate and amino acids as gluconeogenic substrates, and lipolysis of adipose tissue and the release of NEFAs that are taken up by hepatocytes and can be used for β-oxidation or re-esterified and stored as TG. The rate of NEFA re-esterification typically is greater than β-oxidation resulting in transient steatosis after fasting and with insulin resistance or other metabolic dysfunction contributes to NAFLD development^28,29^. Since *Ppp1r3b*^*∆hep*^ mice have rapid fasting hypoglycemia while *Ppp1r3b*^*hepOE*^ mice maintain normal blood glucose levels with extended fasting^7^, we examined how changes in *Ppp1r3b* expression affected circulating NEFAs. In *ad lib* fed mice there was no difference in plasma NEFAs among the three groups (**Fig. 3a**). However, *Ppp1r3b*^*∆hep*^ mice fasted for 4 hours have significantly elevated plasma NEFAs compared to *Ppp1r3b*^*f/f*^ and *Ppp1r3b*^*hepOE*^ mice (*Ppp1r3b*^*f/f*^: 0.4 meq/L; *Ppp1r3b*^*∆hep*^: 0.6 meq/L; *Ppp1r3b*^*hepOE*^: 0.4 meq/L; **Fig. 3b**). Liver mRNA expression of *Cd36*, the primary transporter of NEFAs in hepatocytes that has been demonstrated to contribute to steatosis^30^, had a trend towards increased expression in *Ppp1r3b*^*∆hep*^ mice, while *Ppp1r3b*^*hepOE*^ mice has significantly decreased expression (**Fig. 3c**), suggesting *Ppp1r3b*^*hepOE*^ mice have could have decreased uptake of adipose-derived NEFAs that confers additional protection from steatosis.

**Figure 3.**
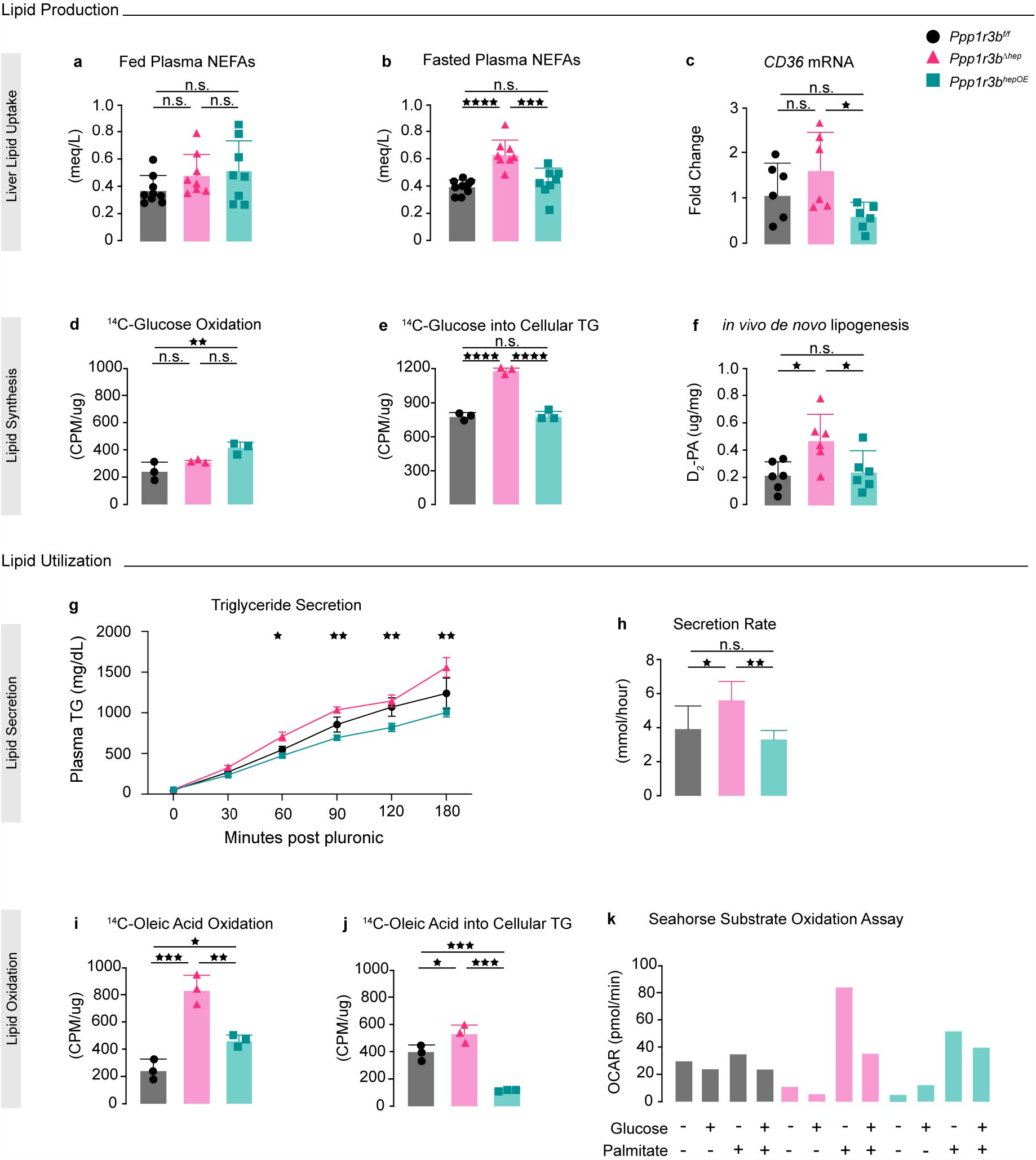
Changes in hepatocyte Ppp1r3b expression alter glucose and lipid metabolism. Young (8 weeks old) mice characterized for metabolic changes in response to hepatic deletion or overexpression of *Ppp1r3b*. <u>Top</u>: Lipid accumulation pathways: **a**. *ad libitum* fed plasma NEFAs, **b**. Plasma NEFAs after 4 hours fasting (*Ppp1r3b*^*f/f*^ (black, n=9), *Ppp1r3b*^*∆hep*^ (pink, n=8), and *Ppp1r3b*^*hepOE*^ (green, n=8) mice. **c**. Liver mRNA expression levels of lipid transporter *Cd36* in *Ppp1r3b*^*f/f*^ (black, n=6), *Ppp1r3b*^*∆hep*^ (pink, n=6), and *Ppp1r3b*^*hepOE*^ (green, n=6) mice. **d**. Oxidation of ^14^C-glucose measured in primary hepatocytes isolated from mice fasted overnight (16 hours). **e**. Incorporation of ^14^C-glucose from media into cellular TG in primary hepatocytes. **f**. *in vivo de novo* lipogenesis assessed by deuterated water incorporation into newly synthesized liver palmitate in *Ppp1r3b*^*f/f*^ (black, n=6), *Ppp1r3b*^*∆hep*^ (pink, n=6), and *Ppp1r3b*^*hepOE*^ (green, n=6) mice. <u>Bottom</u>: Lipid utilization pathways: **g**. in vivo TG secretion as total plasma concentration and **h**. rate of secretion in *Ppp1r3b*^*f/f*^ (black, n=5), *Ppp1r3b*^*∆hep*^ (pink, n=5), and *Ppp1r3b*^*hepOE*^ (green, n=6) mice. **i**. Oxidation of ^14^C-oleic acid in primary hepatocytes isolated from mice fasted overnight (16 hours). **j**. Incorporation of ^14^C-oleic acid from media into cellular TG in primary hepatocytes. **k**. Seahorse metabolic substrate oxidation assay of primary hepatocytes isolated from fasted *Ppp1r3b*^*f/f*^ (black, n=3), *Ppp1r3b*^*∆hep*^ (pink, n=3), and *Ppp1r3b*^*hepOE*^ (green, n=3) mice, tested in either starvation media or supplemented with glucose, palmitate, or both. Oxygen consumption rate (OCR) graphed as an indication of substrate utilization for ATP production. Group average presented as bar + s.d. with individual animals represented as shapes.

In normal liver, postprandial glucose is stored as glycogen for a source of rapid glucose production during short-term fasting or other energy demands. *Ppp1r3b*^*∆hep*^ mice are unable to store glucose as hepatic glycogen, and we hypothesized that hepatic glucose may be shunted to the DNL pathway resulting in steatosis. We previously reported that *Ppp1r3b*^*∆hep*^ mice do not exhibit glycogenic compensation by other tissues including skeletal muscle or kidney^7^, and therefore we traced the fate of exogenous glucose in hepatocytes with altered *Ppp1r3b* expression. Primary hepatocytes isolated from 8-weeks-old overnight fasted female mice were labeled with ^14^C-glucose (1µCi/mL) for two hours then chased in the label-free media for two hours, then the chase media were assayed for oxidized glucose by measuring radiolabeled CO_2_, and cellular lipids were extracted and measured for radiolabeled TG to determine the amount of labeled glucose stored as lipid^31^. Hepatocytes from *Ppp1r3b*^*hepOE*^ mice oxidized more glucose than *Ppp1r3b*^*f/f*^ hepatocytes, but there was no difference in the amount of oxidized glucose in *Ppp1r3b*^*∆hep*^ hepatocytes (**Fig. 3d**). Interestingly, *Ppp1r3b*^*∆hep*^ hepatocytes had a dramatic increase in the amount of ^14^C-glucose that was stored as cellular TG (**Fig. 3e, Supp. Fig 1**). We then assessed *in vivo* DNL by administering deuterated water (D_2_O) to overnight fasted mice that were refed for three hours and then measured newly synthesized palmitic acid (D_2_-PA) in the livers. Consistent with the ex vivo results, the *Ppp1r3b*^*∆hep*^ mice had a significant increase in hepatic D_2_-PA compared to both *Ppp1r3b*^*f/f*^ and *Ppp1r3b*^*hepOE*^ mice (*Ppp1r3b*^*f/f*^: 0.22 ug/mg tissue; *Ppp1r3b*^*∆hep*^: 0.48 ug/mg; *Ppp1r3b*^*hepOE*^: 0.2 ug/mg; **Fig. 3f**).

Even with the changes in glucose utilization and DNL observed in *Ppp1r3b*^*∆hep*^ mice, differences in liver TG were not detected in 8-weeks-old mice (**Fig. 2m**), so we next investigated lipid utilization pathways. Hepatocytes clear excess lipids from the liver by synthesizing TG and secreting them in VLDL particles, so we tested if hepatic TG clearance is altered by *Ppp1r3b* expression. TG secretion was measured in fasted 8-weeks-old mice, and *Ppp1r3b*^*∆hep*^ mice displayed increased plasma TG as soon as 1 hour after the start of the assay, and TG were secreted from the liver at a faster rate than *Ppp1r3b*^*f/f*^ or *Ppp1r3b*^*hepOE*^ mice (**Fig. 3g-h**), suggesting *Ppp1r3b*^*∆hep*^ mice have greater hepatic TG clearance during fasting, possibly due to the lack of hepatic glycogen. Blood metabolites were also measured during this assay and as expected, *Ppp1r3b*^*∆hep*^ mice had significantly lower blood glucose levels throughout the experiment (Supp. Fig. 2a), consistent with our prior reports of hypoglycemia after four or more hours of fasting. There were no changes in blood lactate levels in any of the groups, but *Ppp1r3b*^*∆hep*^ mice had increased blood ketones that were amplified with fasting duration (Supp. Fig. 2b-c), suggesting *Ppp1r3b*^*∆hep*^ mice rely on ketogenesis during fasting to produce energetic substrates. Elevated blood ketones result from β-oxidation of lipids, another lipid clearance mechanism influencing steatosis. We measured the rate of β-oxidation in primary hepatocytes isolated from mice fasted overnight by labeling cells with ^14^C-oleic acid. *Ppp1r3b*^*∆hep*^ hepatocytes had an increased rate of lipid oxidation when maintained in low glucose (5.5mM) media (**Fig. 3i**). Hepatocytes from *Ppp1r3b*^*∆hep*^ mice also had increased incorporation of the exogenous ^14^C-oleic acid into cellular lipids (**Fig. 3j**). Although lower than *Ppp1r3b*^*∆hep*^ hepatocytes, *Ppp1r3b*^*hepOE*^ cells had increased β-oxidation compared to *Ppp1r3b*^*f/f*^ cells (**Fig. 3i**) and had very low incorporation of exogenous lipids into cellular triglycerides (**Fig. 3j**). These data support previous findings that the increased liver glycogen in *Ppp1r3b*^*hepOE*^ mice is sufficient to supply glucose during fasting and therefore the mice are less reliant on β-oxidation for energy. Surprisingly, *Ppp1r3b*^*∆hep*^ cells had increased β-oxidation and cellular TG accumulation simultaneously, indicating the presence of a futile cycle of lipid storage and utilization in *Ppp1r3b*^*∆hep*^ mice. Such futile cycles have been previously described as an early protective adaptation in response to increased hepatic lipid accumulation that may wane with increasing metabolic dysfunction and liver damage^32^. Increased ketogenesis may metabolize as much as two-thirds of hepatic lipids^32^, which could explain the lack of steatosis in young, chow-fed *Ppp1r3b*^*∆hep*^ mice whereas older or HSD-challenged mice have increased liver lipids. These results suggest that although *Ppp1r3b*^*∆hep*^ mice have increased DNL, they also have compensatory changes in hepatic TG utilization during fasting that confer protection against steatosis in young mice with normal feeding conditions. We investigated this possibility in metabolic substrate utilization assays by Seahorse XF Substrate Oxidation Stress Test. Primary hepatocytes isolated from overnight fasted (16 hours) mice were assayed for glucose and palmitate oxidation preference. Hepatocytes from *Ppp1r3b*^*f/f*^ mice had similar maximal mitochondrial respiration rates when media was supplemented with glucose (5 mM), palmitic acid (160 µM), or both (**Fig. 3k**). *Ppp1r3b*^*∆hep*^ hepatocytes preferentially oxidized palmitic acid and while *Ppp1r3b*^*hepOE*^ hepatocytes did oxidize exogenous lipids, they more efficiently oxidized glucose (**Fig. 3k**). The lipid utilization preference observed with *Ppp1r3b* deletion was further supported by improved metabolic profiles in *Ppp1r3b*^*∆hep*^ mice fed a high fat diet (45% lard) for 12 weeks. *Ppp1r3b*^*∆hep*^ mice did not develop insulin resistance and did not have increased hepatic TG compared to *Ppp1r3b*^*f/f*^ controls (Supp. Fig 3), as opposed to the severe insulin resistance and steatosis phenotypes observed in *Ppp1r3b*^*∆hep*^ mice on HSD (**Fig. 1**).

Taken together, these findings clearly demonstrate that changes in hepatocyte *Ppp1r3b* expression effectively cause a switch between lipid and glucose utilization as energetic substrates. This switch in metabolic efficiencies appears rapidly after changes in *Ppp1r3b* expression, as the metabolic changes are measurable after only 3 weeks of altered hepatic *Ppp1r3b* expression. It is intriguing that *Ppp1r3b*^*∆hep*^ mice have a dramatic adaptive response to impaired glycogenesis by storing exogenous glucose as hepatic TG, which is then efficiently oxidized and secreted upon fasting to compensate for the lack of hepatic glycogen. These metabolic changes are present even without differences in total tissue TG accumulation in *Ppp1r3b*^*∆hep*^ mice, and the pericentral lipid droplet accumulation recapitulates early steatosis phenotypes seen in humans.

Given the results from our mouse models and the replicated associations of altered *PPP1R3B* expression with dysglycemia, plasma lipids, and liver traits in human GWAS, we questioned if indications of these metabolic dysregulations are present in human carriers of genetic variants at the *PPP1R3B* locus. To clarify the effects of changes in *PPP1R3B* expression on hepatic fat and plasma ALT, we interrogated the Penn Medicine BioBank (PMBB), a large, ethnically diverse biorepository with whole-exome sequencing data, plasma ALT values, and hepatic fat quantifications derived from clinical CT scans using a machine learning approach, for carriers of noncoding variants associated with increased *PPP1R3B* expression and for putative loss-of-function (pLOF) *PPP1R3B* variants. Consistent with published findings, carriers of rs4240624, rs4841132, and rs9987289 (all in strong LD) had elevated ALT but *decreased* hepatic fat estimated by CT attenuation (**Fig. 4a**). Carriers of *PPP1R3B* pLOF variants were very rare in this dataset (n=14) which has also been reported by other investigators^15^ and in the Genome Aggregation Database (gnomAD) (Supp. Fig. 4). Heterozygous pLOF variant carriers of European ancestry in PMBB had significantly higher CT-derived hepatic fat (β= 20.83; p= 0.0012), and analysis of the rare pLOF variants by gene burden analysis revealed an increased risk of clinical NAFLD diagnosis (OR= 8.175, p= 7.8e^-5^) (Supp. Fig. 5). We next identified participants from the UK Biobank who were carriers of either rs4841132-A or *PPP1R3B* pLOF variants and who had plasma metabolomics data available. Heterozygous carriers of rs4841132-A had elevated plasma glucose and significantly reduced plasma lipids and ketones compared to noncarriers, matching the traits and directionality of prior GWAS associations and our mouse data (**Fig. 4b**). Heterozygous carriers of *PPP1R3B* pLOF variants exhibited elevated plasma ketones and lipids compared to noncarriers, although the data did not reach significance due to low carrier frequency (**Fig. 4c**).

**Figure 4.**
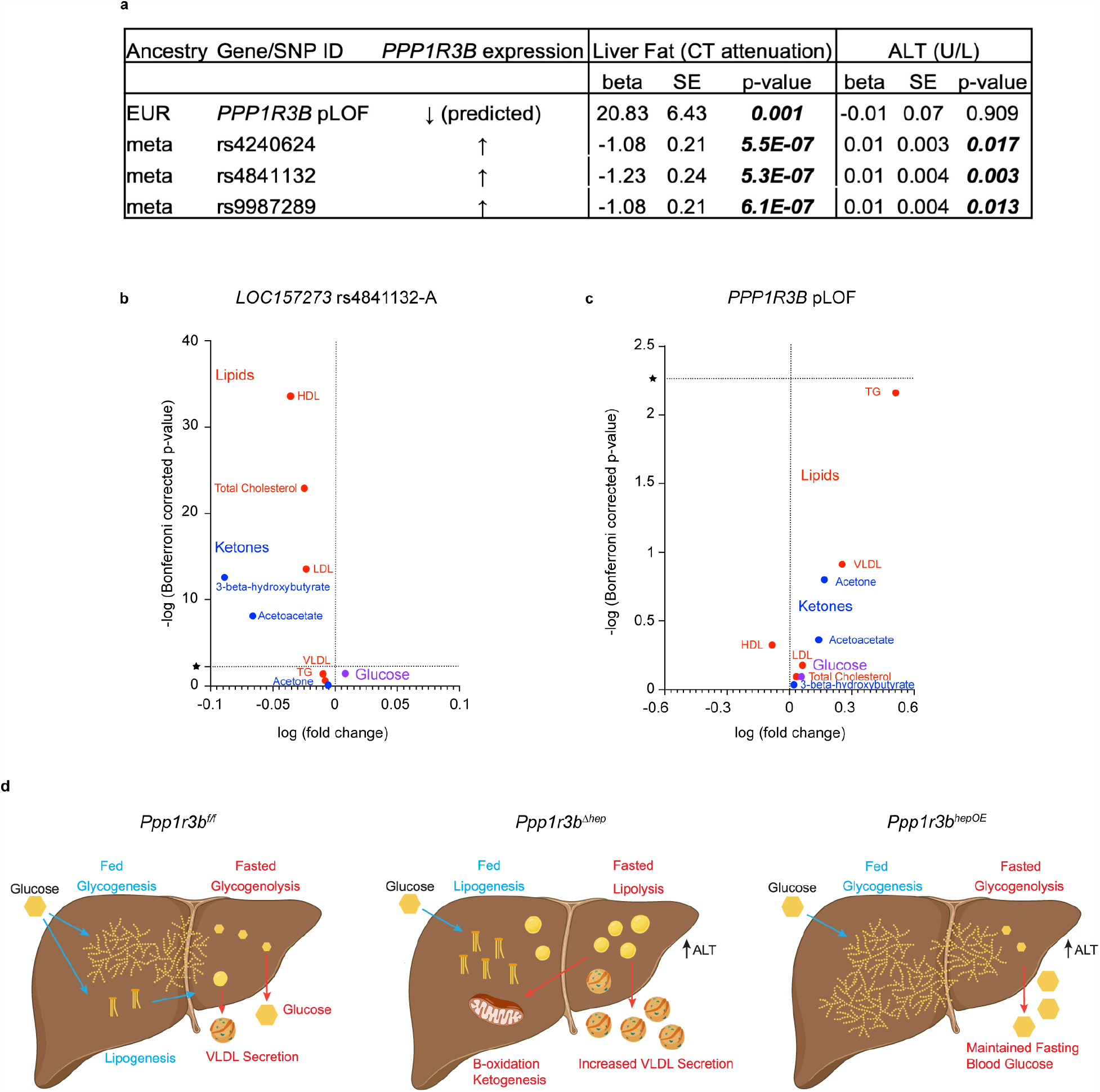
Characteristics of *LOC157273/PPP1R3B* variant carriers corroborate mouse *Ppp1r3b*^*∆hep*^ and *Ppp1r3b*^*hepOE*^ phenotypes. **a**. Analysis of Penn Medicine Biobank participants with Whole Exome Sequencing and CT-derived hepatic fat quantitation yielded a small cohort of individuals with heterozygous *PPP1R3B* pLOF mutations (n=14) as well as carriers of common SNPs in *LOC157273* (n=41,754). Beta indicates the direction of effect (>0 indicates increased trait metric, <0 indicates decreased trait metric), SE =standard error, significant p-values indicated by bold italic font. **b**. Plasma metabolomics data from UK Biobank participants with *LOC157273* rs4841132-A SNPs (left, n=17) and *PPP1R3B* pLOF variants (right, n=25). Horizontal line indicates Bonferroni-corrected p-value with significant metabolites above the line. Vertical line indicates direction of metabolite plasma concentration(>0 indicates increased metabolite, <0 indicates decreased metabolite). **c**. Model of the effects of altered *Ppp1r3b* expression on liver metabolism. <u>Left</u>: with wildtype *Ppp1r3b* expression, postprandial glucose is mainly stored as hepatic glycogen with excess glucose stored as TG. In early fasting, glycogen is converted to free glucose to maintain blood glucose levels and meet extrahepatic energy demands, with lipid utilization and TG secretion as VLDL when glycogen is low. <u>Center</u>: with deletion of hepatocyte *Ppp1r3b* expression, postprandial glycogen synthesis is impaired and exogenous glucose is shunted to lipogenesis and stored as hepatic TG. During fasting, stored TG is oxidized to produce ketones, and lipid is secreted as VLDL. <u>Right</u>: overexpression of hepatocyte *Ppp1r3b* permits enhanced glycogenesis with increased glycogen stores to maintain blood glucose during fasting.

The *PPP1R3B/LOC157273* locus has been associated with glycemic traits, plasma lipids, and liver traits including elevated plasma ALT and hepatic fat. The encoded protein PPP1R3B is a modulator of hepatic storage of glucose as glycogen with an established molecular role contributing to the glycemic GWAS traits but had unclear mechanistic links to liver lipid metabolism and damage phenotypes. Here, we present compelling data demonstrating the role of PPP1R3B in driving hepatic glucose fate in the liver and revealing complex metabolic responses to perturbations in the glycogen metabolism pathway as a result of dysregulated hepatic *Ppp1r3b* expression. With wildtype *Ppp1r3b* expression, postprandial glucose is stored as hepatic glycogen through glycogenesis (**Fig. 4d**, left). Upon fasting, glycogenolysis liberates glucose molecules to be exported from the liver to maintain blood glucose levels. Deletion of hepatocyte *Ppp1r3b* in mice results in the inability to store exogenous glucose as glycogen, leading to fasting hypoglycemia and glucose intolerance. During feeding, *Ppp1r3b*^*∆hep*^ mice shunt glucose to lipogenesis and store both exogenous glucose and lipids as hepatic TG. *Ppp1r3b*^*∆hep*^ mice appear to rapidly adapt to the lack of glycogen by substituting hepatic TG as a fasting energy source with increased β-oxidation and ketogenesis (**Fig. 4d**, center). However, with age or HSD, lipid production and storage exceed lipid utilization resulting in steatosis and liver damage indicated by elevated plasma ALT. The reciprocal phenotype is seen in *Ppp1r3b*^*hepOE*^ mice, in which increased liver glycogen stores maintain blood glucose during fasting and glycogen/glucose is the preferred metabolic substrate even with prolonged fasting. *Ppp1r3b*^*hepOE*^ mice are protected from hepatic steatosis and do not have altered lipid metabolism in response to fasting or refeeding (**Fig. 4d**, right). Consequently, the increased hepatic glycogen stores in *Ppp1r3b*^*hepOE*^ mice may contributes to glycogenosis with elevated plasma ALT, recapitulating the human phenotypes associated with GWAS SNPs at this locus. Intriguingly, plasma metabolomics in human carriers of rare pLOF mutation in *PPP1R3B* are concordant with the lipid energy preference seen in *Ppp1r3b*^*∆hep*^ mice, further supporting that PPP1R3B is critical in regulating hepatic glucose storage and consequential energy supply during fasting. These studies provide novel insights into the role of PPP1R3B as a metabolic switch between hepatic glycogen and lipid storage and demonstrate that changes in *Ppp1r3b* expression, either by deletion or overexpression, contribute to altered glucose metabolism, liver damage, and cardiometabolic disease risk.

## Methods

### Generation of Hepatocyte-Specific Ppp1r3b Knockout and Overexpressing Mice

Mice with hepatocyte-specific deletion (*Ppp1r3b*^*∆hep*^) and overexpression (*Ppp1r3b*^*hepOE*^) were achieved as previously described^7^. Briefly, *Ppp1r3b*^*f/f*^ mice on C57BL/6J background produced for Merck by Taconic were bred to mice expressing Cre recombinase driven by the Albumin promoter (*Alb-Cre*), with littermate mice without *Alb-Cre* transgene expression used as controls. *Ppp1r3b* overexpression was induced by administering adeno-associated virus serotype 8 with thyroxine binding globulin promoter-driven murine *Ppp1r3b* (AAV8.TBG.Ppp1r3b; titer 1e^12^ gc) and littermates were administered AAV with an empty expression cassette as control (AAV8.TBG.Null; titer 1e^12^ gc). In some experiments, *Ppp1r3b*^*f/f*^ mice were administered AAV with TBG promoter-driven Cre recombinase expression (AAV8.TBG.Cre; titer 1.5e^11^ gc) to generate hepatocyte-specific *Ppp1r3b* knockout mice. Littermates were administered either AAV8.TBG.Ppp1r3b for overexpression or AAV8.TBG.Null as control (both titers 1.5e^11^ gc). AAV vectors were generated by the University of Pennsylvania Vector Core (Philadelphia, PA, USA). Both male and female mice were assessed unless specifically indicated.

### Animal Housing and Diet

Animals were housed under controlled temperature (23°C) and lighting (12-hour light/dark cycle) with free access to water and standard mouse chow (LabDiet 5010), or animals were administered High Sucrose Diet (66% Sucrose Content, Research Diets D11725i) or High Fat Diet (45% lard, Research Diets D12451i) for 12 weeks. All animal experiments were reviewed and approved by Institutional Animal Care and Use Committee and the University Laboratory Animal Resources Committee of the University of Pennsylvania, Philadelphia, USA.

### In vivo Metabolic Testing

For glucose tolerance tests (GTT), mice were fasted overnight (14-16h) then administered 2 g/kg fasted body mass glucose by intraperitoneal (i.p.) injection with subsequent measurements of veinous tail blood glucose with a OneTouch Ultra Mini glucometer (LifeScan) at the indicated timepoints. Insulin tolerance tests (ITT) were performed in mice after 6 hours fasting by i.p. administration of recombinant insulin (Novolin R; Novodisk A/S; 0.75 U/kg body mass) and subsequent measurement of veinous tail blood glucose with a OneTouch Ultra Mini glucometer (LifeScan) at the indicated timepoints.

### In vivo de Novo Lipogenesis

Mice were fasted overnight (16 hours), then weighed and a baseline blood collection was obtained from retro orbital bleeding. Mice were administered deuterated water (Sigma-Aldrich; 400 µL/20 g body mass) and allowed to feed *ad libitum* for 3 hours. Mice were then sacrificed and approximately 100 mg liver tissue and 50 µl plasma were provided to the Institute of Diabetes, Obesity and Metabolism (IDOM) Metabolic Tracer Resource at the University of Pennsylvania for analysis of body water and palmitate deuterium enrichment by gas chromatography-electron impact ionization mass spectrometry and subsequent calculation of *de novo* lipogenesis^33^.

### In vivo TG Secretion

Hepatic TG secretion was assessed as previously described^34^. Male mice were fasted for 4 hours then administered Pluronic P407 (30mg in 200uL 1xPBS) retro-orbitally, then bled at the indicated timepoints. Plasma TG content was measured colorimetrically using Infinity Triglyceride Reagent (ThermoScientific) according to the manufacturer’s protocol. Hepatic TG secretion rates were calculated from the slope of the regression line of the time versus plasma TG.

### Blood and plasma metabolites

Blood glucose, lactate and ketone bodies were measured in veinous tail blood by OneTouch Ultra Mini glucometer (LifeScan), Lactate Plus (Nova Biomedical), or novaMax Plus (Nova Biomedical) meters, respectively. Plasma insulin was measured by ELISA (ChrystalChem Cat#90080). Plasma non-esterified fatty acids (NEFA) and alanine aminotransferase (ALT) were measured by ACE Alfa Wasserman Axcel autoanalyzer using commercially available colorimetric assay kits (Wako).

#### Metabolic and Biochemical Measurements

Hepatic TG were measured colorimetrically using Infinity Triglyceride Reagent (ThermoScientific). Tissues were homogenized in PBS, diluted 1:5, and incubated with 1% deoxycholate then assayed according to the manufacturer protocol. Sample TG was calculated by standard curve and normalized to wet tissue mass. Hepatic glycogen levels were measured by Glycogen Assay Kit II (Colorimetric) from Abcam (Cat# ab169558). Tissues were homogenized with distilled H2O on ice and then boiled for 10 min. Homogenates were spun at 13,000 r.p.m. for 10 min and supernatants were assayed for glycogen content. Results were normalized by protein content.

### Histology and Immunohistochemistry

Tissue sections were paraffin-embedded, sectioned, and stained by the Histology and Gene Expression Co-Op at Penn Cardiovascular Institute. Standard techniques were applied to stain tissues with hematoxylin and eosin (H&E) to assess general tissue morphology, Periodic acid-Schiff (PAS) to detect polysaccharide content, and Picro-Sirius Red to detect collagen deposition and liver lesions. Immunohistochemistry (IHC) was performed on de-paraffinized sections to detect lipid droplets with PLIN2 antibodies (Sigma-Aldrich HPA016607). Brightfield images were captured on a slide scanning microscope (Keyence BZ-X800) and are displayed at 4x resolution.

### RNA Isolation and Quantitative RT-PCR

Total RNA was extracted from approximately 100mg flash-frozen liver tissue following homogenization in RNAzol RT reagent (Sigma-Aldrich). cDNA was synthesized from 1ug RNA using High-Capacity cDNA Reverse Transcription Kit (Applied Biosystems). Quantitative real-time PCR (qPCR) was performed with SYBR Green Mix (Applied Biosystems) on an QuantStudio™ 7 Flex Real-Time PCR System (Thermo Fisher). Oligonucleotide primer sequences for target and reference genes are listed in Supplemental Table 1. The fold change in the target mRNA abundance with respect to the control group and normalized by the reference genes was calculated using the 2^-∆∆C^_T_ method.

### Primary Hepatocyte Isolation and ex vivo Oxidation Assays

Primary hepatocytes were isolated from overnight fasted (16 hours) female mice by a two-step perfusion with collagenase digestion as described previously^35^. Primary hepatocytes were seeded at a density of 5×10^5^ cells/well in triplicate wells of 6-well plates. After 2 hours attachment in glucose-free DMEM, cells were washed and labeled with either ^14^C-oleic acid (1µCi/mL) for β -oxidation or ^14^C-glucose (1µCi/mL) for glucose oxidation^31,36^. After 2 hours, label media were removed, cells gently washed, and chased for 2 hours. Chase media were transferred to sealed oxidation flasks and treated with 70% perchloric acid with gentle rocking for 1 hour to capture ^14^C-CO_2_ on KOH-soaked filter paper. The filter paper and incompletely oxidized intermediates (acid-soluble metabolites) were counted for radioactivity. Cells were washed and lipids extracted by 3:2 hexane:isopropanol, then dried and resuspended in 100% hexane for separation by thin layer chromatography, and the TG bands were cut and counted for radioactivity. The cells were then washed and cellular protein solubilized and quantified by BCA to normalize the radioactivity measurements for each well.

### Measurement of Substrate Oxidation by Seahorse Analysis

The Seahorse Bioscience XF96 Extracellular Flux analyzer (Agilent) was used to quantify mitochondrial respiration of primary hepatocytes in the presence of nutrient-free media or media supplemented with glucose, palmitic acid, or both. Cells were seeded at 12×10^3 cells/well in 96-well XF96 plates and incubated in nutrient-free media for 1 hour. After cells adhered, media were gently aspirated and replaced with either nutrient-free media, glucose media (5 mM glucose), lipid media (160 µM BSA-Palmitic acid), or combined glucose and lipid media. Cells were incubated for 4 hours then gently washed, and Seahorse Assay Media added (180 µL/well) and incubated at 37° without CO_2_ for 45 min. The substrate oxidation assay was performed with automated addition of ATP synthase inhibitor (1.5 µM oligomycin), proton uncoupler (2 µM FCCP), and electron transport chain inhibitors (0.5 µM rotenone/antimycin A mixture). The oxygen consumption rates (OCR) were measured and analyzed using Seahorse Wave software.

### Analysis of LOC157273 SNPs and PPP1R3B predicted loss-of-function variants in Penn Medicine Biobank

All individuals recruited for the Penn Medicine BioBank (PMBB) are patients of clinical practice sites of the University of Pennsylvania Health System. This study was approved by the Institutional Review Board of the University of Pennsylvania and complied with the principles set out in the Declaration of Helsinki. PMBB participants who had both whole exome sequence data and CT-derived hepatic fat quantitation (n=10,283) were analyzed for carriage of the *LOC157273* SNPs rs4240624, rs4841132, and rs9987289, or for rare (minor allele frequency <0.1% in gnomAD) potential loss-of-function (pLOF) variants and rare predicted deleterious (REVEL > 0.8) missense (pDM) variants in *PPP1R3B*, in the genetic region chr8:8997073-9009084 of GRCH38^37,38^. Analysis of each of the *LOC157273* SNPs and a gene burden of rare pLOF and pDM variants in *PPP1R3B* were associated with hepatic fat using a linear regression model adjusted for age, genetically determined sex, and principal components (PC) of ancestry (PC1-5 in Africans, PC1-10 in Europeans). We used an additive genetic model to aggregate variants as previously described^39^. These analyses were performed separately by African and European genetic ancestry and combined with inverse variance weighted meta-analysis.

### Phenome-wide association of hepatic fat with EHR diagnoses and traits

A phenome-wide association study (PheWAS) was performed to determine the phenotypes associated with the quantitative trait of median hepatic fat in PMBB for the 10,283 unrelated individuals in PMBB with both exome sequences and quantitated hepatic fat available^40^. ICD-10 encounter diagnoses were mapped to ICD-9 via the Center for Medicare and Medicaid Services 2017 General Equivalency Mappings (https://www.cms.gov/Medicare/Coding/ICD10/2017-ICD-10-CM-and-GEMs.html) and manual curation. Phenotypes for each individual were then determined by mapping ICD-9 codes to distinct disease entities (*i*.*e*., Phecodes) using the R package “PheWAS”^41^. Patients were determined to have a certain disease phenotype if they had the corresponding ICD diagnosis on 2 or more dates, while phenotypic controls consisted of individuals who never had the ICD code. Individuals with an ICD diagnosis on only one date as well as individuals under control exclusion criteria based on PheWAS phenotype mapping protocols were not considered in statistical analyses. Each Phecode was tested for association with quantitated hepatic fat using a logistic regression model adjusted for age, sex, and principal components (PC1-10) of genetic ancestry. The association analyses considered only disease phenotypes with at least 20 cases based on power calculations in a prior simulation study^39^. This led to the interrogation of 1396 total Phecodes, and we used a Bonferroni correction to adjust for multiple testing (p= 0.05/1396= 3.58e^-05^).

### Plasma metabolomics in UK Biobank participants

The UK biobank is a population-based cohort study conducted in the United Kingdom from 2006 to 2010, which recruited 502,500 volunteers aged 37 to 73 years at baseline. All participants were registered with the UK National Health Service and attended an initial examination, which is followed by a long-term follow-up taking place continuously. On the baseline visit, blood samples were taken, and biometric measures were performed. In a subgroup of UKB participants (n= 105,348) NMR-based plasma metabolomic profiling was performed. Only participants with genetic information and metabolomic profiling were included in the analyses. Plasma glucose, ketone bodies, and major lipid classes were chosen *a priori* for further analysis. Each metabolite was divided by its standard deviation and calculated the log fold change between variant carriers and controls as well as a -log transformed logistic regression. Differences were considered to be statistically significant when p<0.05. Bonferroni correction was performed to avoid first type of error occurring through multiple testing of n=9 metabolites (p<0.05/9= 0.0056). The data were analyzed using R version 4.0.2 (R Foundation for Statistical Computing; Vienna, Austria), SPSS Statistics version 26 (IBM; Armonk, NY, USA) and Prism version 8 (GraphPad, LaJolla, CA, USA).

### Statistical Analyses

All results are presented as mean +S.D unless otherwise indicated. Data are presented as one representative experiment and each experiment was repeated 2-3 times with consistent outcomes. Results were analyzed by the unpaired two-tailed Student’s t-test or one-way ANOVA as appropriate using GraphPad Prism Software (Version 9.3.1, GraphPad Software, Inc.). Statistical significance was defined as *p<0.05, **p<0.01, ***p<0.001, ****p<0001.

## Acknowledgments

This research has been conducted using the UK Biobank Resource under Application Number 70653. We thank the University of Pennsylvania Diabetes Research Center (DRC) for the use of the Radioimmunoassay and Biomarkers Core and Metabolic Tracer Core (P30-DK19525), and the Cardiovascular Institute Histology and Gene Expression Co-Op for histological services.

This work was supported by NIH R01DK114291 (K.T.C., N.J.H, J.A.B., D.J.R.), UM1DK126194 (K.T.C. and D.J.R.).

