## Supplemental figures for "PPP1R3B is a metabolic switch that shifts hepatic energy storage from lipid to glycogen"

**Figure S1. Thin layer chromatography (TLC) plates of cellular lipids from glucose and β-oxidation studies.** Primary hepatocytes isolated from *Ppp1r3b*$^{f/f}$, *Ppp1r3b*$^{\Delta hep}$, and *Ppp1r3b*$^{hepOE}$ were assayed for metabolism of $^{14}$C-labeled glucose (left) or oleic acid (right). Cellular lipids were extracted and separated by TLC. The TG bands were cut and counted for radioactivity and normalized to cellular protein.

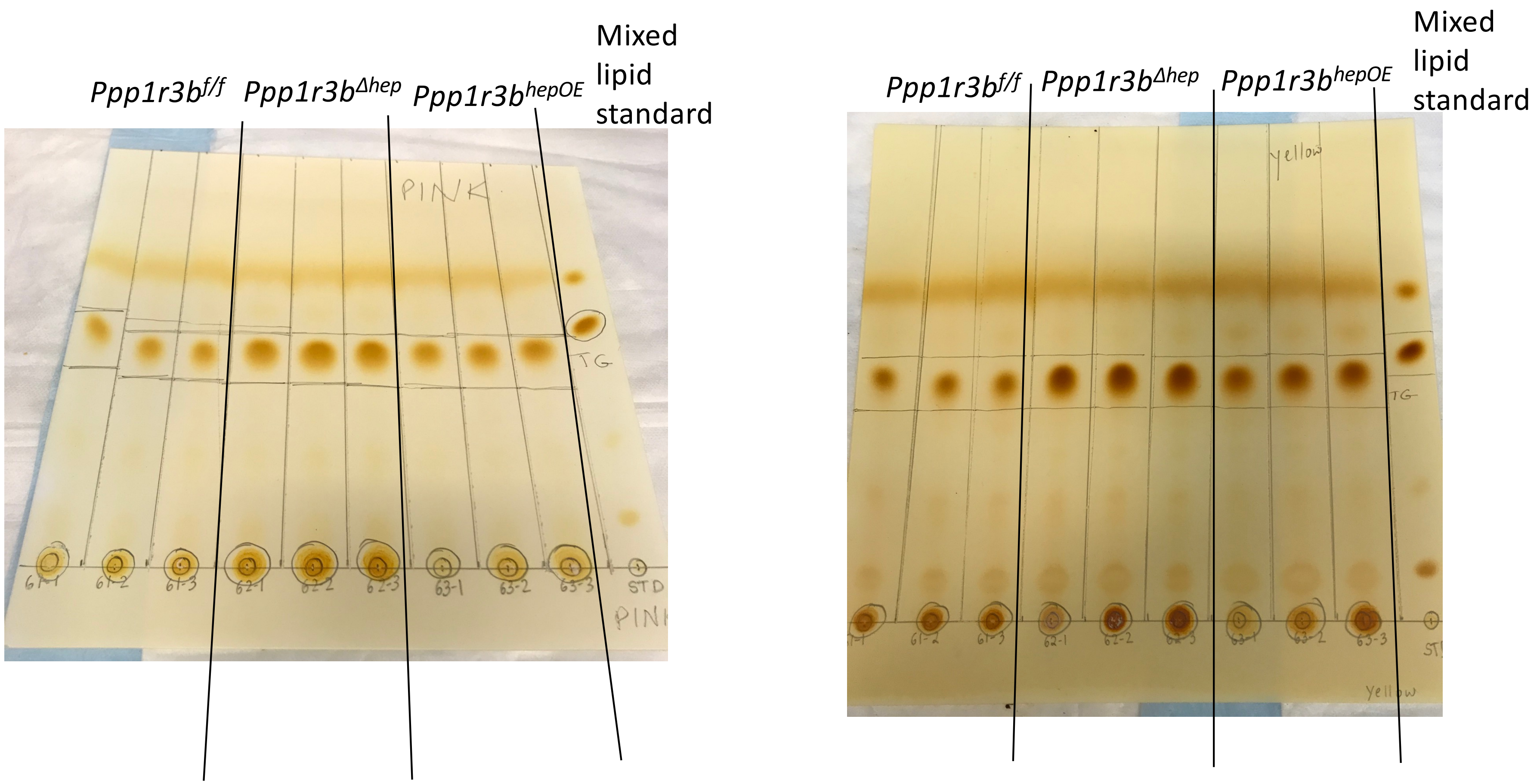

**Figure S2. Fasting blood metabolites during TG secretion assay.** Mice were fasted for 4 hours then administered pluronic-P407 to inhibit peripheral lipolysis during in vivo TG secretion assays. Fasting blood levels of glucose (**a**), lactate (**b**), and ketones (**c**) were measured at the indicated timepoints.

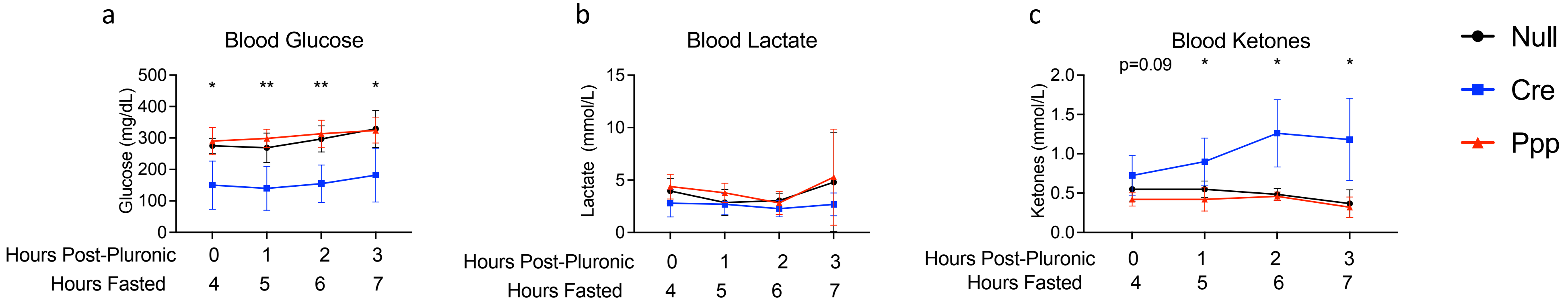

**Figure S3. *Ppp1r3b$^{\Delta hep}$* mice do not develop insulin resistance or exhibit increased steatosis on HFD.** *Ppp1r3b$^{f/f}$* and *Ppp1r3b$^{\Delta hep}$* mice were fed HFD for 12 weeks. **a.** Glucose tolerance test after 8 weeks of HFD feeding. **b.** Insulin tolerance test after 10 weeks HFD. **c.** Liver glycogen and **d.** liver TG measured after 12 weeks HFD.

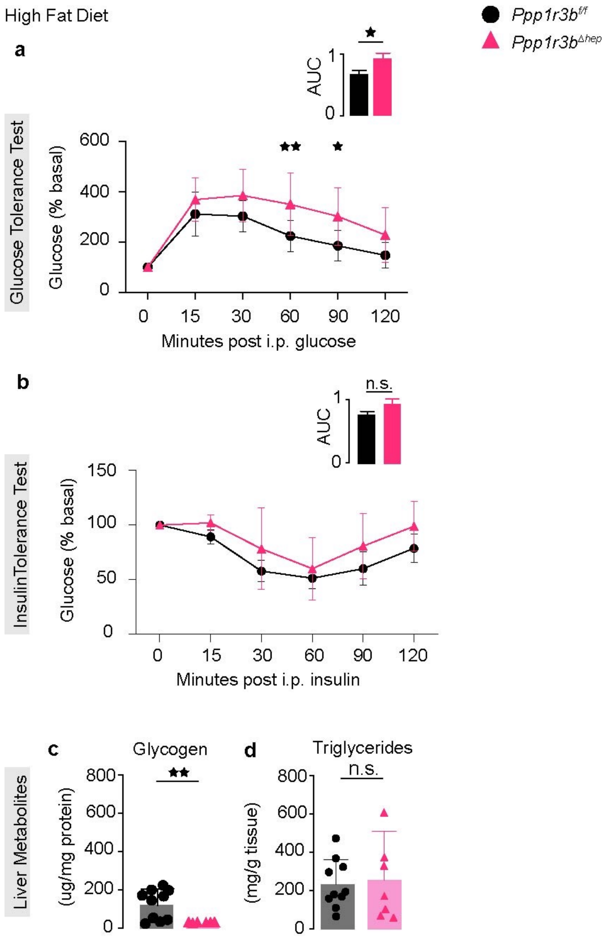

**Figure S4. Genetic variation in the *PPP1R3B* region reported in gnomAD indicates fewer than expected pLOF mutations.** Observed synonymous and missense mutations in *PPP1R3B* are within expected frequencies, but pLOF mutations are reported significantly less than expected. Screenshot of database search performed 15-Oct-2022, accessible at https://gnomad.broadinstitute.org/gene/ENSG00000173281?dataset=gnomad_r2_1

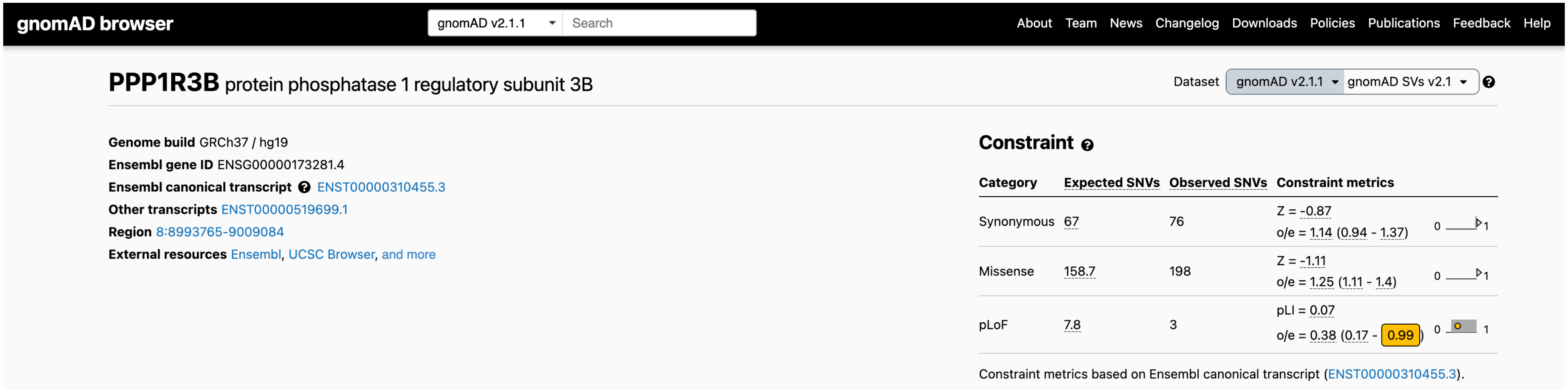

**Figure S5. Phenome-wide associations of rare *PPP1R3B* pLOF variants.** Ancestry-specific phenotypes associated with PPP1R3B pLOF variants in PMBB. AFR= African ancestry, EUR= European, Meta= AFR + EUR.

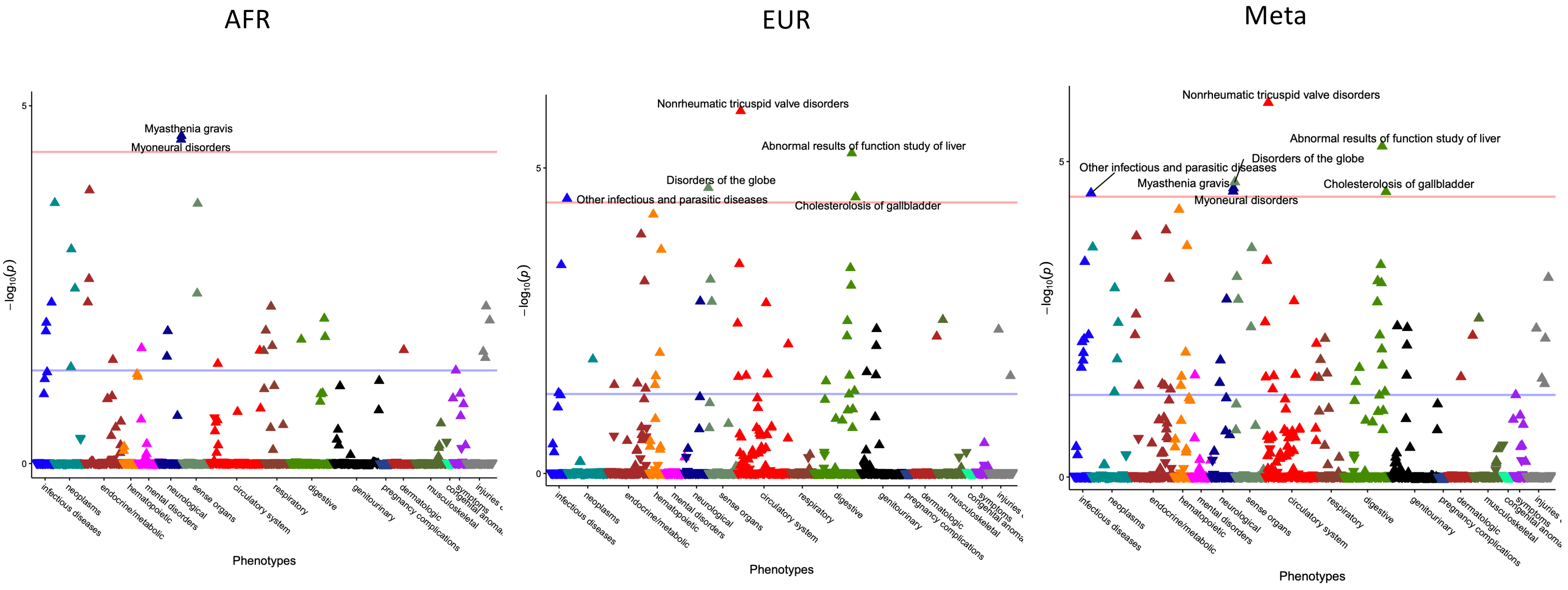

**Table S1. *PPP1R3B* pLOF variants in PMBB.** Summary of variant locations and number of carriers of rare pLOF variants in PMBB used for gene burden analysis.

carrier_freq

| | variant | n_0 | n_1 | n_2 |
|---|---|---|---|---|
| 1 | 8:9140905:D:1_T | 41758 | 1 | NA |
| 2 | 8:9140972:C:A_A | 41758 | 1 | NA |
| 3 | 8:9141077:A:G_G | 41758 | 1 | NA |
| 4 | 8:9141117:A:C_C | 41757 | 1 | NA |
| 5 | 8:9141126:D:1_G | 41757 | 2 | NA |
| 6 | 8:9141158:G:A_A | 41757 | 2 | NA |
| 7 | 8:9141174:C:T_T | 41758 | 1 | NA |
| 8 | 8:9141180:T:C_C | 41758 | 1 | NA |
| 9 | 8:9141186:A:T_T | 41758 | 1 | NA |
| 10 | 8:9141212:C:T_T | 41758 | 1 | NA |
| 11 | 8:9141594:D:2_C | 41758 | 1 | NA |
| 12 | 8:9141600:I:1_GC | 41754 | 1 | NA |
